# Throat color polymorphism is related to differences in aggression in the Aegean wall lizard

**DOI:** 10.1101/2024.03.14.585063

**Authors:** Dhruthi S. Mandavilli, Ian J. Wang, Kinsey M. Brock

## Abstract

Aggressive behavior can be used to establish and maintain access to crucial resources such as space, food, and mating opportunities. Color polymorphic animals sometimes exhibit morph-correlated aggressive behaviors that can influence relative reproductive success and, thus, the maintenance of polymorphism. The Aegean wall lizard, *Podarcis erhardii*, exhibits three monochromatic throat color morphs: orange, white, and yellow. Previously, male *P. erhardii* color morphs were shown to differ in their use of aggressive behaviors and ability to win staged contests during laboratory experiments. However, whether these color morphs use aggressive behavior differently in their natural setting where ecological and environmental factors are not standardized remains unknown. Here, we used *in situ* observations of wall lizard color morph interactions over a large section of dry stone wall to investigate behavioral differences in aggression among color morphs *in situ*. We compared the counts and intensities (aggression scores) of aggressive behaviors (both performing and receiving aggression) among color morphs and found that color morphs differ significantly in the frequencies and intensities of their aggressive behaviors. We found that the white morph exhibited significantly more aggression than the orange and yellow morphs on dry stone walls. Taken together, results from *in* and *ex situ* behavioral studies suggest that the smaller, more common white color morphs are more aggressive, which might help explain their relatively greater abundance and persistence across the species’ range.

---

LIZARDS use aggressive behaviors to access crucial resources such as food, space, and mating opportunities (Stamps, 1977; Andersson, 1994; Taborsky et al. 2008; Hurtado-Gonzales and Uy, 2010). This aggressive behavior occurs when animals send threatening signals or engage in physical combat. Lizards signal aggression in many ways, including flashing colorful parts of the body (Thompson and Moore, 1991), making themselves appear larger (Robson and Miles, 2000), gaping (Brock et al., 2022a), and performing head bobs and tail displays (Robson and Miles, 2000; Peters et al., 2016). Physical aggression in lizards ranges in intensity from behaviors like chasing and lunging to mounting, wrestling, and biting (Stamps, 1977).

Aggression can be influenced by hormone levels (Greenberg and Crews, 1990; Moore and Thompson, 1990), population adult sex ratios (Le Galliard et al., 2005), population density (Donihue et al., 2016), resource availability (Wu et al., 2019), and social or behavioral strategies (McLean et al., 2015; Yewers et al., 2016; Brock et al., 2022a).

Alternative behavioral strategies associated with genetically-based color polymorphisms are common in lizards (Sinervo and Lively, 1996; Robson and Miles, 2000; Stuart-Fox et al., 2020; Brock et al., 2022a). Some lizard color morphs with alternative behavioral strategies exhibit different levels of aggression (Bastiaans et al., 2013; Abalos et al., 2016; Yewers et al., 2016; Brock et al., 2022a), and certain color morphs even exhibit morph aggression bias where they adjust their aggressive behaviors depending on the morph with which they are interacting (Sinervo and Lively 1996; Yewers et al., 2016; Scali et al., 2021). Because more aggression in lizards is associated with greater access to resources and greater reproductive success (Herrel et al., 2009), differential morph aggression could play important roles in the evolution and maintenance of color polymorphism (Sinervo and Lively, 1996; Dijkstra et al., 2008; Horton et al., 2012; Yewers et al., 2016; Tinghitella et al., 2018; Scali et al., 2021). Using aggressive behaviors to win contests over limited resources such as space and mating opportunities can increase morph relative fitness and reproductive success and, in turn, affect morph allele frequencies (Sinervo and Lively, 1996; Tinghitella et al., 2018). If morphs of the same color compete more often or exhibit like-morph aggression bias then the color morph at the lowest frequency in the population would experience less aggression than more common morphs (Seehausen and Schluter, 2004), thereby creating social conditions that may allow the least common morph to increase in frequency in the population, a type of Allee effect (Kingston et al., 2003; Qvarnström et al., 2012; Dhiman and Poria, 2018). Thus, morph aggressive interactions can play a role in balancing selection via negative frequency-dependent selection that maintains morph diversity over long timescales (Sinervo and Lively, 1996; Seehausen and Schluter, 2004; Scali et al., 2021).

Most *Podarcis* wall lizard species are color polymorphic and have genetically-determined orange, white, and yellow ventral color morphs (Andrade et al., 2019; Brock et al., 2022b). The Aegean wall lizard, *Podarcis erhardii*, is color polymorphic and exhibits morph differences in social, competitive, and anti-predator behaviors (Brock and Madden, 2022; Brock et al., 2022a). Male *P. erhardii* color morphs also differ in body size and maximum bite force (Brock et al. 2020). Male *P. erhardii* orange morphs are larger and have a stronger bite force (due to a larger head size) than white and yellow males (Brock et al., 2020). Larger body size and red-orange coloration often correlate with more testosterone, higher levels of aggression, and physical and social dominance in birds and reptiles (Tokarz, 1985; Pryke, 2009; Olsson et al., 2013).

However, male *P. erhardii* color morphs exhibited the opposite pattern in *ex situ* behavioral trials conducted under standardized laboratory conditions where lizards were size-matched and neither lizard had a residency advantage (Brock et al., 2022a). In these laboratory experiments, orange morphs performed significantly fewer aggressive behaviors than white and yellow morphs while competing over limited basking space on a dry stone wall in the lab. White morphs, which tend to be smaller (Brock et al., 2020), exhibited the highest levels of aggression and delivered the most bites during these laboratory contests (Brock et al., 2022a). White and yellow morphs were associated with winning, and orange morphs were associated with losing contests over thermally desirable basking space (Brock et al., 2022a). Laboratory behavioral experiments allow us to eliminate the influence of certain variables (e.g. size-matching lizards and neutrality of the arena to control the influence of body size and home-field advantage). However, lizards are not size-matched in nature and might adjust their aggressiveness based on an opponent’s size and other habitat and residency factors (Stamps, 1977; Stuart-Fox and Johnston, 2005; Sacchi et al., 2009). Thus, observations of natural interactions between color morphs *in situ* are necessary to more fully understand the role of aggression in morph social dynamics and alternative behavioral strategies.

In this study, we used *in situ* observations of *P. erhardii* color morph interactions to determine whether morph aggression levels differ under natural conditions. We counted aggressive behaviors during one-on-one interactions to test if morphs perform aggressive behaviors such as gaping, chasing, lunging, and biting at different frequencies when in their natural habitat. We then assigned points to these aggressive behaviors to determine whether color morphs perform and receive different amounts of aggression. Finally, we counted the number of conspecific bite scars on 127 lizards from the study area to test whether certain morphs experience more overall aggression independent of sex and body size. We expected that the larger orange morphs would perform more non-physical aggressive behavior, such as gaping, to signal their size advantage and potentially avoid expending energy on more vigorous behaviors, like fighting. Given that white and yellow morphs tend to be smaller (Brock et al., 2020), we expected that these morphs would perform physically aggressive behaviors more often than the orange morph to access space, food, and mating opportunities on the dry stone walls. Because physical aggressive behaviors are more intense than signaling aggressive behaviors, we expected that white and yellow morphs would have higher aggression scores than orange morphs. Finally, we predicted that white morphs would have more conspecific bite scars than other morphs because they are smaller and easier to bite and seem to instigate higher-intensity aggressive interactions more often than the other morphs (Brock et al., 2022a).

## Methods

### Study Species

*Podarcis erhardii* is a small to medium-sized lacertid lizard (adult snout-vent-length, SVL, 45-80 mm) that occurs in a variety of habitats across the southern Balkans, including many Aegean islands (Gruber, 1987; Valakos et al., 2008; Speybroeck et al., 2016). True to its common name, this wall lizard is often found on dry stone walls that provide a range of thermally suitable basking spaces (Fig 1). *P. erhardii* is diurnal and has a bimodal activity period during the hot summer months characterized by a longer morning activity period (0800-1200 hrs), followed by a shorter evening activity period (1600-1900 hrs; Valakos, 1990). The *P. erhardii* breeding season peaks from April to June (Edsman, 2001), during which *P. erhardii* exhibits high levels of social activity and frequently engages in aggressive behaviors such as biting, chasing, gaping, and lunging (Brock et al., 2022a).

**FIG 1:**
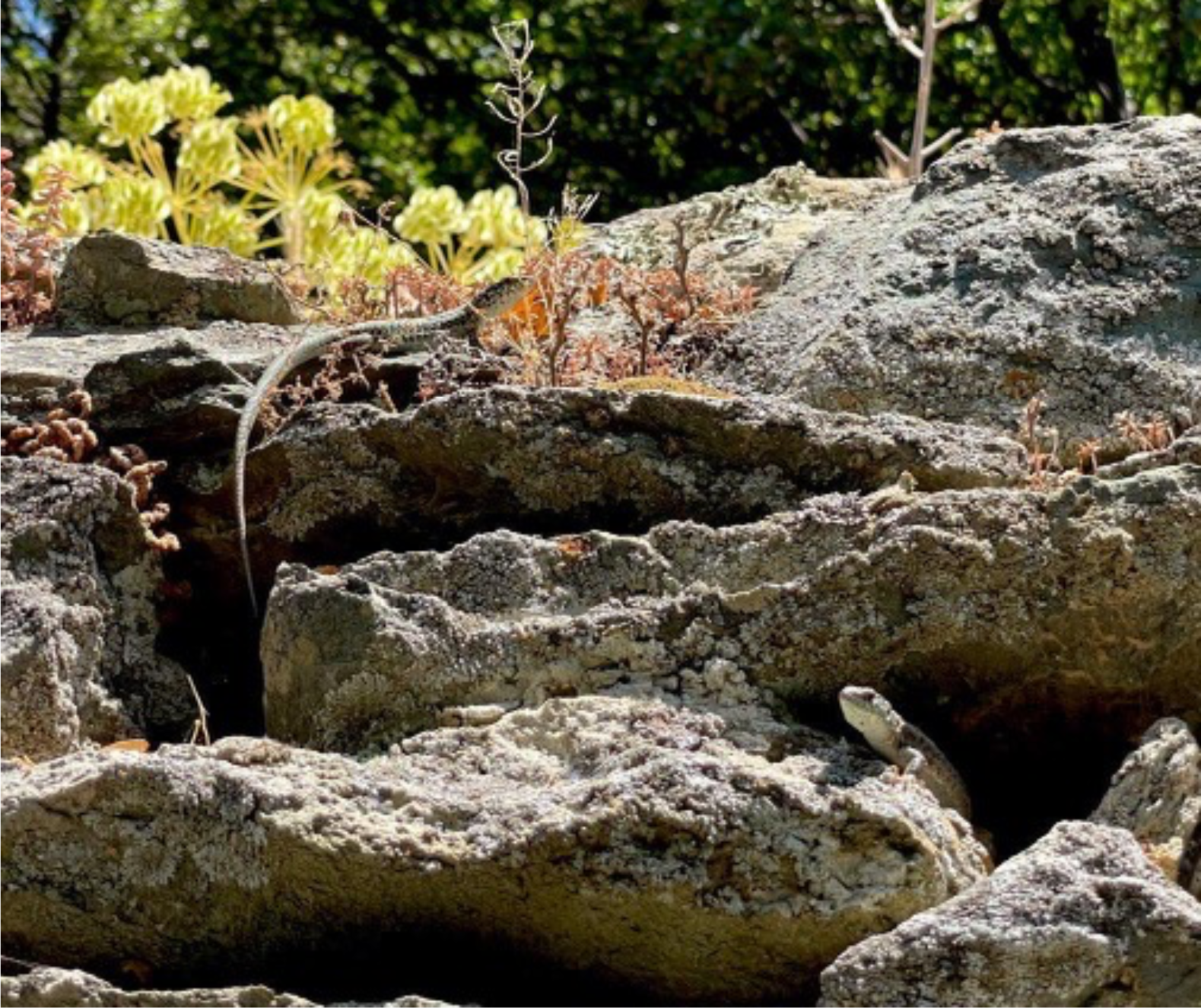
Two *P. erhardii* white morphs inspecting each other on a dry stone wall moments before an aggressive interaction.

In *P. erhardii* aggression toward conspecifics usually starts with non-physical visual signals like throat signaling (lifting the head back to expose the throat to the aggressee), dorsolateral flattening with side displays, and gaping (opening one’s mouth wide toward the aggressee; Brock et al., 2022a). Aggressive encounters can escalate quickly to physical altercations, such as lunging (hitting the aggressee anywhere on the body using one’s head in a fast, forward motion) and chasing (quickly pursuing the aggressee). The most intense form of physically aggressive behavior *P. erhardii* exhibits is biting (using one’s mouth to clamp down on the aggressee), which can cause wounds, leave scars, and even kill (Vervust et al., 2009; Cooper et al., 2015; Donihue et al., 2016; Madden and Brock, 2018).

### *In Situ* Behavioral Observations

We conducted our study on the island of Naxos, Greece, in July 2022. Naxos is the largest Cycladic island located in the center of *P. erhardii*’s island distribution and has all three monochromatic color morphs (Brock et al., 2022c). We observed lizards from a single population in the agricultural village of Moni (37.081694° N, 25.493056° E). Moni has large sections of dry stone walls mixed with low, spiny vegetation and olive trees. These stone walls host a dense population of *P. erhardii* that can be observed easily at a distance without being disturbed. We chose this study site for its abundance of lizards and to compare our results to a previous behavioral study from the same population that was conducted in a laboratory setting (Brock et al., 2022a).

We delineated an 8 m long by 2.5 m high section of stone wall for our study area with flagging tape. We chose this section of the dry stone wall for its seclusion from passersby and high density of lizards. To observe lizards without influencing their behavior, we sat behind an olive tree (approximately 5.5 m away from the center of the stone wall study area), viewed the study area through binoculars (Nikon Monarch M5), and made observations for three days during *P. erhardii’s* peak daily activity period (0800-1300). When we observed two lizards facing each other we considered that the beginning of a social interaction. Once a social interaction began, we noted the color and sex of the lizards and observed these individuals until they disengaged from the interaction. We considered looking or walking away as disengagement. If aggression (i.e. gaping, chasing, lunging, or biting) occurred, we noted which lizard performed the aggressive behavior (the “aggressor”) and who the aggression was directed toward (the “aggressee”).

### Aggression Scores

To determine which morphs exhibit higher levels of aggression we calculated an aggression score for every interaction we observed. Following other studies on wall lizards, we used a point system that reflects the intensity of the various aggressive behaviors (Abalos et al., 2016; Brock et al., 2022a). Biting, the most aggressive behavior, was given the highest point value of 4, followed by lunging (3 points), chasing (2 points), and gaping (1 point). Thus, a higher aggression score indicates higher overall levels of aggression. We calculated aggression scores for aggressors in each interaction by summing the total number of points per interaction. To calculate a score for the average level of aggression each morph receives, we averaged the sums of aggression scores directed toward all of the individuals within each morph.

### Bite Scar Counts

We counted the number of conspecific bite scars over the entire body on 127 lizards we captured from Moni to assess color morph differences in receiving high intensity aggression outside of our observational period. Intraspecific bite scars are easily distinguishable from scars produced by predators due to their shape and size, so we disregarded any scarring not clearly inflicted by a conspecific (Donihue et al., 2016). We identified lizards as female if they had reduced, inactive femoral pores and male if they had large, active femoral pores. Finally, we measured the SVL of every captured lizard with a pair of digital precision calipers (Mitutoyo 500-171-30 Absolute Scale Digital Caliper, Aurora, Illinois, USA). We returned each lizard to its point of capture after these measurements.

### Ethical Statement

The methods used in this research were approved by the Greek Ministry of Energy and the Environment (permit ΥΠΕΝ/ΔΔΔ/5619/145) and the University of California, Berkeley Institutional Animal Care and Use Committee (protocol AUP-2021-08-14567).

### Analyses

We used RStudio (v 4.2.1; R Core Team, 2022) for all statistical analyses and data visualization. To determine whether color morphs differentially perform aggressive behaviors (gaping, chasing, lunging, and biting) in their natural habitat, we used chi-square goodness of fit tests and subsequent post-hoc tests: pairwise *t*-tests with a Bonferroni correction, comparing the number of observed occurrences per aggressive behavior between each color morph using the functions ‘chisq.test’ and ‘pairwise.t.test’ in the ‘stats’ package in R (v 4.2.1; R Core Team, 2022). Next, we tested for differences in morph aggressor and aggressee scores calculated from each interaction using ANOVA or Kruskal-Wallis tests. We first ran Shapiro-Wilk and Levene’s tests using the ‘shapiro.test’ and ‘leveneTest’ functions from the ‘stats’ and ‘car’ packages to check that aggression score data met assumptions for ANOVA testing of normality and variance, respectively (v 4.2.1; R Core Team, 2022; v 3.1.0; Fox and Weisberg, 2019). When the assumptions for ANOVA were not met, we ran the nonparametric Kruskal-Wallis and post-hoc Dunn tests using the ‘kruskal.test’ and ‘dunnTest’ functions from the ‘stats’ and ‘FSA’ packages (v 4.2.1; R Core Team, 2022; v 0.9.3; Ogle et al., 2022).

We used multiple linear regression to estimate the effects of morph (factor variable with three levels: orange, white, yellow), sex (character variable with two levels: female, male), and body length (numeric variable: SVL in mm) on conspecific bite scar counts (numeric variable: number of bite scars per individual), implementing the following model structure with the ‘lm’ function in the stats package: bite scars ∼ morph + sex + svl. Lastly, we used chi-square goodness of fit tests and pairwise t-tests with a Bonferroni correction to test for morph-specific differences in conspecific bite scars.

## Results

In total, we observed 34 intermorph and 21 intramorph aggressive interactions (Table 1), and we obtained bite scar counts for 127 lizards in Moni (N = 41 orange lizards, N = 44 white lizards, N = 42 yellow lizards). Of 96 observed social interactions, 55 were aggressive.

**TABLE 1:** Number and type of one-on-one aggressive interactions observed between *P*.

| Aggressor morph | Aggressee morph |  |  |
| --- | --- | --- | --- |
|  | Orange | White | Yellow |
| Orange | 1 | 9 | 3 |
| White | 4 | 14 | 9 |
| Yellow | 2 | 7 | 6 |

We observed different levels of aggressive behaviors among color morphs (Table 2). Morphs differed significantly in how many times they bit other lizards (chi-square test, *χ2* = 29.31, df = 2, *P* < 0.001), with white morphs biting other lizards more times (25 total instances of biting) compared to orange (2 total instances of biting; pairwise t-test, *P* = 0.0034) and yellow morphs (5 total instances of biting; pairwise t-test, *P* = 0.0230). Orange and yellow morphs did not differ significantly in how many times they bit other lizards (pairwise t-test, *P* = 1.0000).

**TABLE 2:** Chi-square goodness of fit test results from morph aggressive behavior counts. White morphs bit, chased, and lunged significantly more than orange and yellow morphs. Morphs did not significantly differ in gaping behavior, thus no *post hoc* test was necessary.

| Behavior | Observed<br>Counts | Expected<br>Counts | Difference<br>(Obs - Exp) | Chi-square<br>goodness of fit | Bonferroni adjustment |
| --- | --- | --- | --- | --- | --- |
| Bite | O = 2 | 10.67 | O = -8.67 | $\chi^2 = 29.31$, df = 2, | O vs W: $P = 0.003$ |
| | W = 25 | | W = 14.33 | $P < 0.001$ | O vs Y: $P = 1$ |
| | Y = 5 | | Y = -5.67 | | W vs Y: $P = 0.02$ |
| Chase | O = 1 | 19.67 | O = -18.67 | $\chi^2 = 80.37$, df = 2, | O vs W: $P < 0.001$ |
| | W = 52 | | W = 32.33 | $P < 0.001$ | O vs Y: $P = 0.47$ |
| | Y = 6 | | Y = -13.67 | | W vs Y: $P < 0.001$ |
| Gape | O = 7 | 10 | O = -3 | $\chi^2 = 5.40$, df = 2, | N/A |
| | W = 16 | | W = 6 | $P = 0.07$ | |
|  | Y = 7 |  | Y = -3 |  |  |
| Lunge | O = 2 | 13 | O = -11 | $\chi^2 = 55.85$, df = 2, | O vs W: $P < 0.001$ |
| | W = 35 | | W = 22 | $P < 0.001$ | O vs Y: $P = 1$ |
| | Y = 2 | | Y = -11 | | W vs Y: $P < 0.001$ |

Additionally, morphs differed significantly in the number of times they chased other lizards (chi-square test, *χ2* = 80.37, df = 2, *P* < 0.001). White morphs chased other lizards far more often (52 times) compared to orange (1 time; pairwise t-test, *P* < 0.001) and yellow morphs (6 times; pairwise t-test, *P* < 0.001). Orange and yellow morphs did not differ significantly in how many times they chased lizards (pairwise t-test, *P* = 0.47). Morphs did not significantly differ in the number of times they gaped at other lizards (chi-square test, *χ2* = 5.4, df = 2, *P* = 0.07).

However, morphs did significantly differ in the number of times they lunged at other lizards (chi-square test, *χ2* = 55.85, df = 2, *P* < 0.001). Once again, white morphs performed this behavior far more often (35 lunges) compared to orange (2 lunges; pairwise t-test, *P* < 0.001) and yellow morphs (2 lunges; pairwise t-test, *P* < 0.001). Orange and yellow morphs did not significantly differ in lunging count (pairwise t-test, *P* = 1).

White morphs displayed higher levels of aggression during intermorph interactions (Fig 2), while all morphs received similar amounts of aggression (Kruskal-Wallis test, *χ2* = 0.08, df = 2, *P* = 0.96; Fig 2). We found that the average aggression scores between orange and white morph aggressors (Kruskal-Wallis test, *χ2* = 32.36, df = 2, *P* < 0.001; Dunn test, *Z* = −4.886, *P*unadj < 0.001, *P*adj < 0.001) and white and yellow morph aggressors (Dunn test, *Z* = 4.40, *P*unadj < 0.001, *P*adj < 0.001) were significantly different. Aggression scores did not differ significantly between orange and yellow morph aggressors (Dunn test, *Z* = −0.61, *P*unadj = 0.54, *P*adj = 0.54).

**FIG 2:**
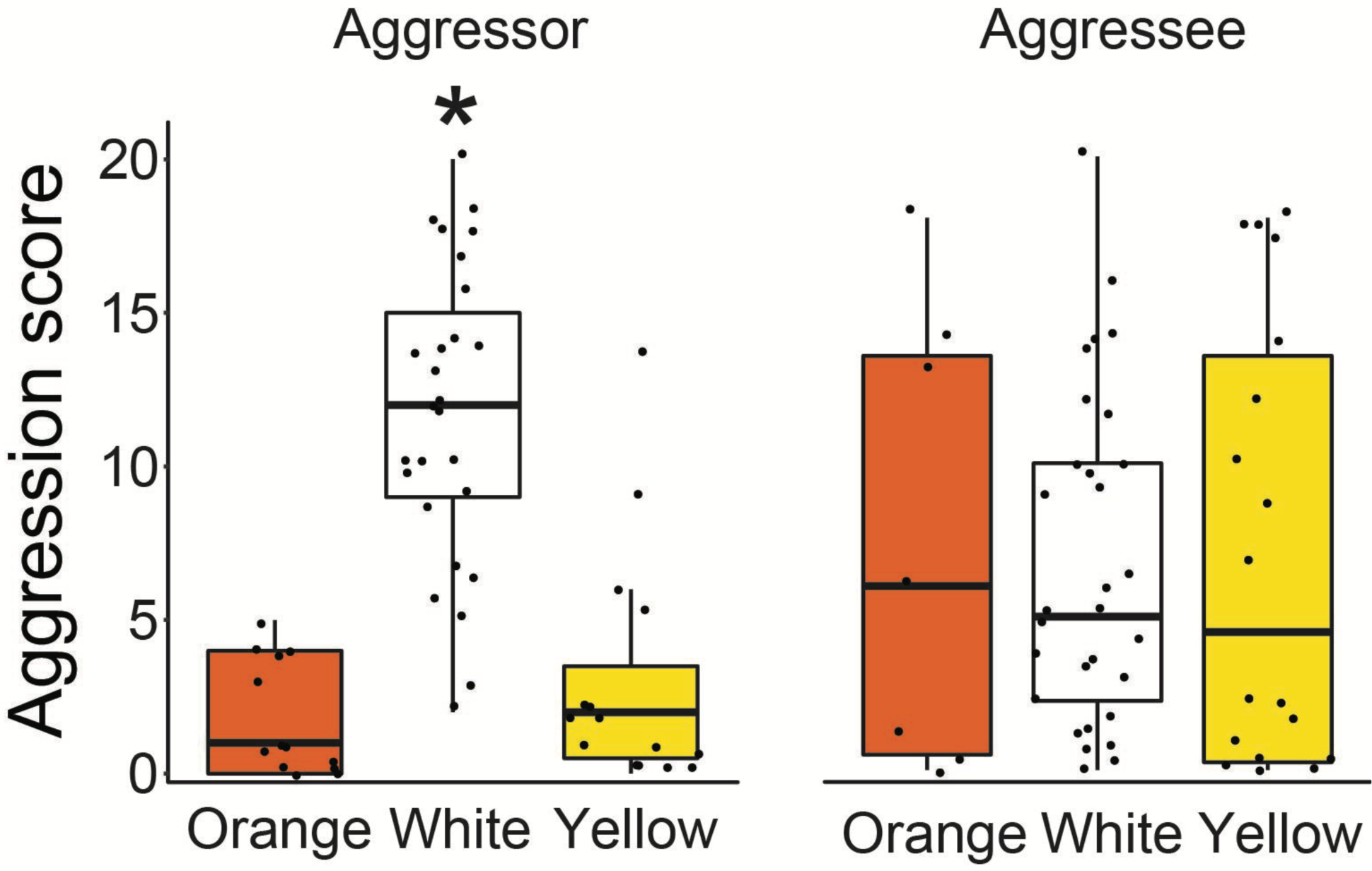
Aggression scores exhibited by (aggressor) and directed toward (aggressee) each *P. erhardii* color morph in box-and-whisker plots, with the whiskers showing minimum and maximum aggression score and boxes representing the first and third quartiles of aggression scores with a median line for each morph. White morphs exhibited significantly higher levels of aggression during intermorph interactions compared to orange and yellow morphs (Dunn test *P* < 0.001, denoted with an asterisk). Morphs did not significantly differ in the level of aggression they received from unlike morphs (Kruskal-Wallis test *P* = 0.96).

Results from the multiple regression model revealed that color morph was a significant predictor of conspecific bite scar count (Table 3). Color morphs had significantly different numbers of bite scars (chi-square test, *χ2* = 66.41, df = 2, *P* < 0.001; Fig 3). Specifically, white morphs exhibited significantly more bite scars than both orange (pairwise *t*-test, *P* = 0.003) and yellow morphs (pairwise *t*-test, *P* < 0.001). Orange and yellow morphs did not have significantly different numbers of conspecific bite scars (pairwise *t*-test, *P* = 0.96).

**FIG 3:**
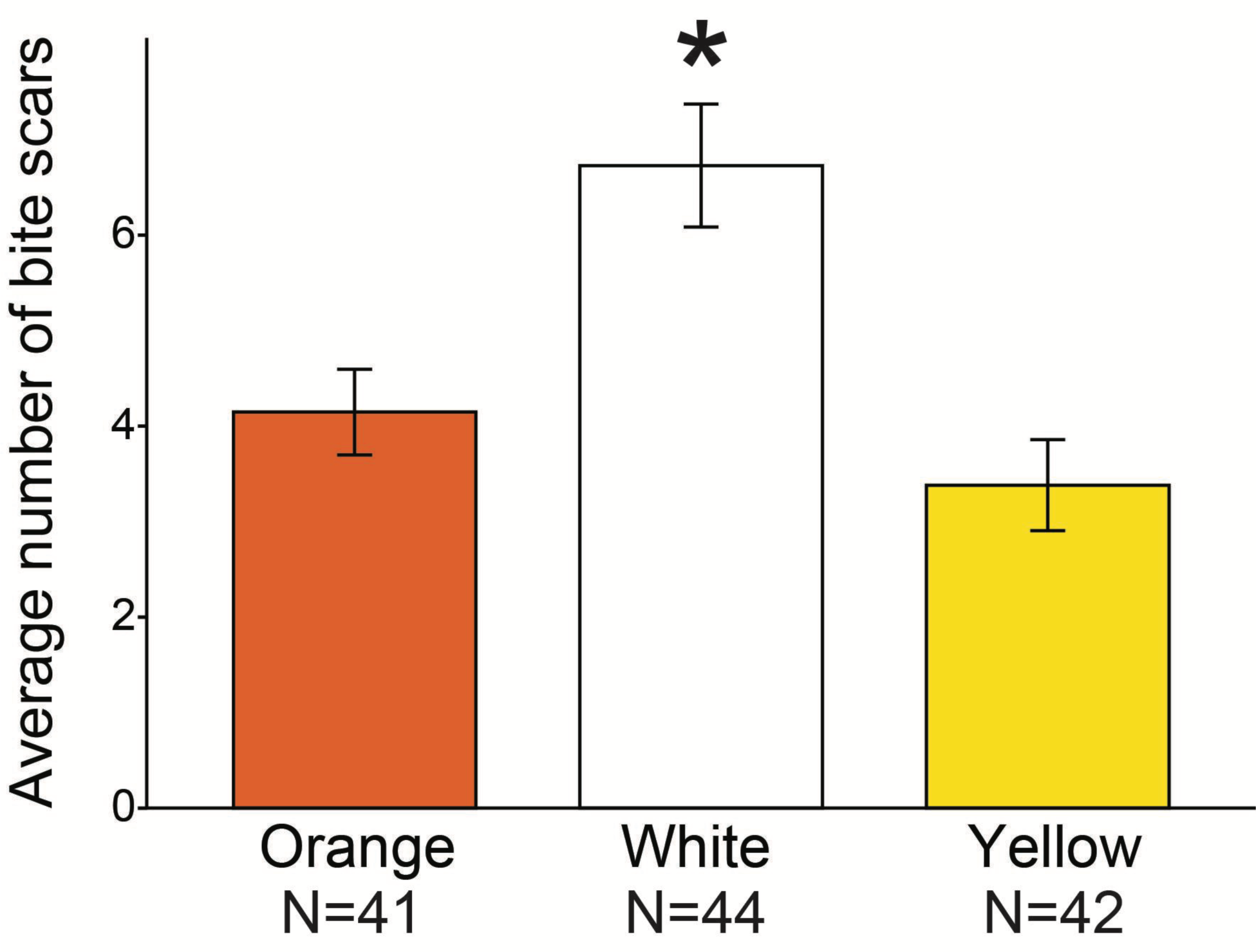
The average number of bite scars per individual on *P. erhardii* color morphs from Moni, Naxos. White morphs (denoted with an asterisk) had significantly more bite scars than both orange (pairwise *t*-test, *P* = 0.003) and yellow morphs (pairwise *t*-test, *P* < 0.001). Sample sizes are noted under each morph.

**TABLE 3:** Bite scar multiple regression model output. The only significant predictor of bite scar count in our dataset was white morphs (*P* < 0.001).

| Predictor | Estimate | Std. error | $t$ value | $P$ value |
| --- | --- | --- | --- | --- |
| White morph | 2.81 | 0.79 | 3.54 | $< 0.001^*$ |
| Yellow morph | -0.82 | 0.79 | -1.04 | 0.30 |
| Sex | -1.27 | 0.70 | -1.82 | 0.07 |
| SVL | 0.14 | 0.07 | 1.91 | 0.06 |

## Discussion

A fundamental goal in biology is to understand how natural variation is generated, maintained, and lost. Color polymorphic taxa exhibit extreme intraspecific phenotypic variation, and the stability and maintenance of that variation over long evolutionary timescales requires some sort of balancing selection among morphs, else the polymorphism is lost. Alternative behavioral strategies are one way that multiple color morphs can persist in a population over long periods (Sinervo and Lively, 1996; Sinervo and Zamudio, 2001; Calsbeek et al., 2010). Aggression is one category of behavior that differs among *P. erhardii* color morphs and could be involved in their alternative behavioral strategies. We found that white morphs performed aggressive behaviors (biting, chasing, and lunging) significantly more often than orange and yellow morphs, corroborating results from a previous laboratory study (Brock et al., 2022a), but all three color morphs experienced similar average levels of aggression directed toward them.

We also found a significant relationship between conspecific bite scar counts and color morph in this species but not with sex or body size, with aggressive white morphs having significantly more bite scars than both the orange and yellow morphs. Since lizard aggression can increase resource access and reproductive success, these higher levels of aggression can enhance fitness and explain differences in color morph frequencies (Taborsky et al. 2008; Herrel et al., 2009; Hurtado-Gonzales and Uy, 2010). Of the lizards we caught in Moni, 75% were white, 18% were yellow, and 7% were orange. Orange morphs are present at relatively low frequencies, sometimes even absent, compared to the other morphs on Naxos and on many other Aegean islands (Brock and Madden, 2022; Brock et al., 2022c). If white morphs are using aggression to win more competitions over territory and mates, then they might have more opportunities to reproduce and, hence, be present at higher frequencies in the population (Coladonato et al., 2020). Theory suggests that alternative strategies have context-dependent advantages and disadvantages to coexist indefinitely (Gross, 1991; Sinervo and Lively, 1996). In several color polymorphic wall lizard species, orange morphs are usually found in cooler, more mesic, highly vegetated areas near water (Capula et al. 2009; I de Lanuza et al., 2018; BeVier et al., 2022; Thompson et al., 2023). Perhaps in the case of *P. erhardii*, smaller, more aggressive white morphs that prefer hotter temperatures have an advantage over orange and yellow morphs in hotter, drier, stone wall habitat, and thus achieve higher fitness and greater frequencies in these environmental contexts at the microhabitat (BeVier et al., 2022; Thompson et al., 2023) and island-level (Brock et al., 2022c).

So, what is the role of differential morph aggression in maintaining the Aegean wall lizard throat color polymorphism? In lizards, color traits often correlate with behavior and can act as important signals to conspecifics (Whiting et al., 2006; I de Lanuza et al., 2014). Even before engaging in physical combat, animals can convey differences in fighting ability using morph color signaling (Arak, 1983; Buinjé et al., 2019). In many species of squamates, carotenoid-based coloration (such as yellow, oranges, and reds) is associated with greater contest performance and competitive ability (Hamilton et al., 2013). Studies of male Augrabies flat lizards (*Platysaurus broadleyi*) and Striped lava lizards (*Tropidosaurus semitaeniatus*) found that individuals with highly-reflective UV throats had higher levels of testosterone, exhibited higher levels of aggression, had greater fighting ability, and obtained more space through aggression (Stapley and Whiting, 2006; Whiting et al., 2006; Bruinjé et al., 2019). A similar signaling phenomenon could be present in *P. erhardii* throat color morphs, as some white morphs have UV reflection on their throats (Brock et al., 2020). Future studies should test if UV-reflectiveness correlates with aggression and fighting ability in *P. erhardii* and if individuals use this as a signal to avoid initiating fights with the white morphs (Brock et al., 2020). Other studies on color polymorphic lizards have found associations between red-orange coloration and higher levels of testosterone, which is broadly associated with aggression in animals (Sinervo et al., 2000; Huyghe et al., 2009; Yewers et al., 2016). In *P. erhardii*, the generally smaller white morphs (some with UV-reflective throats), might need to use aggression to obtain food and mating opportunities in the presence of larger orange and yellow morphs that may more readily access these resources through their larger body size (Brock et al., 2020).

Body and head size are key characteristics that impact contest outcomes in various taxa (Smith and Parker, 1976; Andersson 1994; Olsson and Madsen 1998), and lizards are no exception. Larger lizards usually have higher levels of testosterone and aggression and, thus, tend to win contests over smaller lizards (Tokarz, 1985; Olsson and Shine, 2000; Fairbairn et al., 2007; Arnott and Elwood, 2009; Huyghe et al., 2009). However, Aegean wall lizards did not follow this body size trend in a laboratory contest study conducted on male color morphs (Brock et al., 2022a). In Aegean wall lizards, color morph predicted contest outcomes independent of body size (Brock et al., 2022a). Additionally, the smaller white and yellow morphs won more often and performed more aggressive behaviors during those contests (Brock et al., 2022a). Body size is generally correlated with strength, stamina, and ability to inflict injury in animals (Arnott and Elwood, 2009). In *P. erhardii* the orange morphs tend to have bigger bodies, larger head sizes, and significantly stronger maximum bite force capabilities than the smaller white and yellow morphs (Brock et al., 2020). In some instances, smaller lizards might be more aggressive than larger lizards if aggression is the best way for smaller individuals to gain resources. This phenomenon is known as the “Desperado effect” (Grafen 1987; Schulte-Hostedde and Millar 2002; Just et al. 2007). Since white morphs tend to be smaller (Brock et al., 2020), their stature might not intimidate or ward off physical aggression from other lizards, which could result in more physical altercations and thus more bite scars, as seen in our bite scar results. Indeed, in another study, orange morphs signaled aggression by gaping more often than they used physical aggression in intermorph interactions (Brock et al. 2022a). On the other hand, larger body size can be beneficial when determining contest outcomes without a need for aggression when lizards assess their size differences to see if they are willing to enter a risky physical altercation (Prenter et al. 2008; Yoshino et al. 2011). Correspondingly, we observed that the larger orange morphs had fewer bite scars. Perhaps an orange throat signals larger head and body size, which wards off potential aggressors and results in fewer conspecific bite scars. Yellow morphs exhibit a wide range of body sizes (Brock et al. 2020), tend to use intermediate levels of aggression compared to the orange and white morphs (Brock et al. 2022a), and had relatively few conspecific bite scars. In addition to the orange morph’s intimidating body size, it is possible that orange and yellow morphs avoid getting bitten by conspecifics by choosing microhabitats that are less densely populated. We observed high densities of lizards on the walls, though most of these lizards were white morphs. Wall lizards obtain very high densities in stone wall microhabitat (Boag, 1972), where we conducted our study, whereas orange and yellow tend to be found more often in shaded, moist microhabitat where wall lizards are less densely populated than on walls (BeVier et al. 2022).

Here, we have shown that *P. erhardii* color morphs differ significantly in how often and intensely they express aggressive behavior. White morphs perform aggressive behaviors both more often and intensely than orange and yellow morphs, which could have implications for contest outcomes over limited resources and give white morphs a potential fitness advantage over orange and yellow morphs that is counterbalanced somehow over time. White morphs tend to have more bite scars than the other morphs, which indicate participating in more aggressive interactions to acquire or maintain access to resources such as space, food, and mating opportunities. White morphs were also involved in many more intense aggressive interactions, such as biting (25 instances of biting), especially compared to the orange morph (2 instances of biting). Orange and white morphs in *P. erhardii* seem to be the most distinct from each other in aggression traits, and yellow morphs are intermediate. Along with previous studies (Brock et al. 2022), our results here lend more support to the idea that *P. erhardii* color morphs might have alternative, though not completely discrete, behavioral strategies. All of the color morphs exhibit the same aggressive behaviors, though morphs differ in how often they engage in aggression, which could be due to a combination of genetic and environmental factors. More research on correlations between behavioral traits and fitness outcomes is needed to understand how color polymorphism is maintained in wall lizards across environmental contexts. Future research could observe color morph aggression and contest outcomes in different environmental conditions that favor one morph over the other. For example, we tend to find orange and yellow morphs more often on plant substrates (e.g. trees, wood, spiny shrubs) near shaded areas, with orange morphs often using vegetation for refuge from perceived threats (BeVier et al., 2022; Brock and Madden, 2022). An important and unknown factor to study in many color polymorphic species is how morph behavior and heterogeneous environments together influence the maintenance of morph diversity, especially if morphs have behavioral strategies that impact fitness and are influenced by certain environmental aspects. Color polymorphic wall lizards are an ideal study system in which to study these questions due to similar color morph-environment relationships (e.g., orange morphs associated with wet, cool environments) across *Podarcis* species (Capula et al. 2009; I de Lanuza et al., 2018).

## Acknowledgements

We thank Asher Thompson, Jerry Sun, Tanmayi Patharkar, and Serey Leakna Nouth for their help catching lizards for bite scar counts. This research was funded by a University of California, Berkeley College of Natural Resources Travel Grant and SPUR funds awarded to D.S. Mandavilli and a National Science Foundation Postdoctoral Research Fellowship in Biology (Award Number 2109710) awarded to K.M. Brock.

## Notes

### Competing Interest Statement

The authors have declared no competing interest.

